## Supplemental figure 3 for "Fingerprinting and chemotyping approaches reveal a wide genetic and metabolic diversity among wild hops (*Humulus lupulus* L.)"

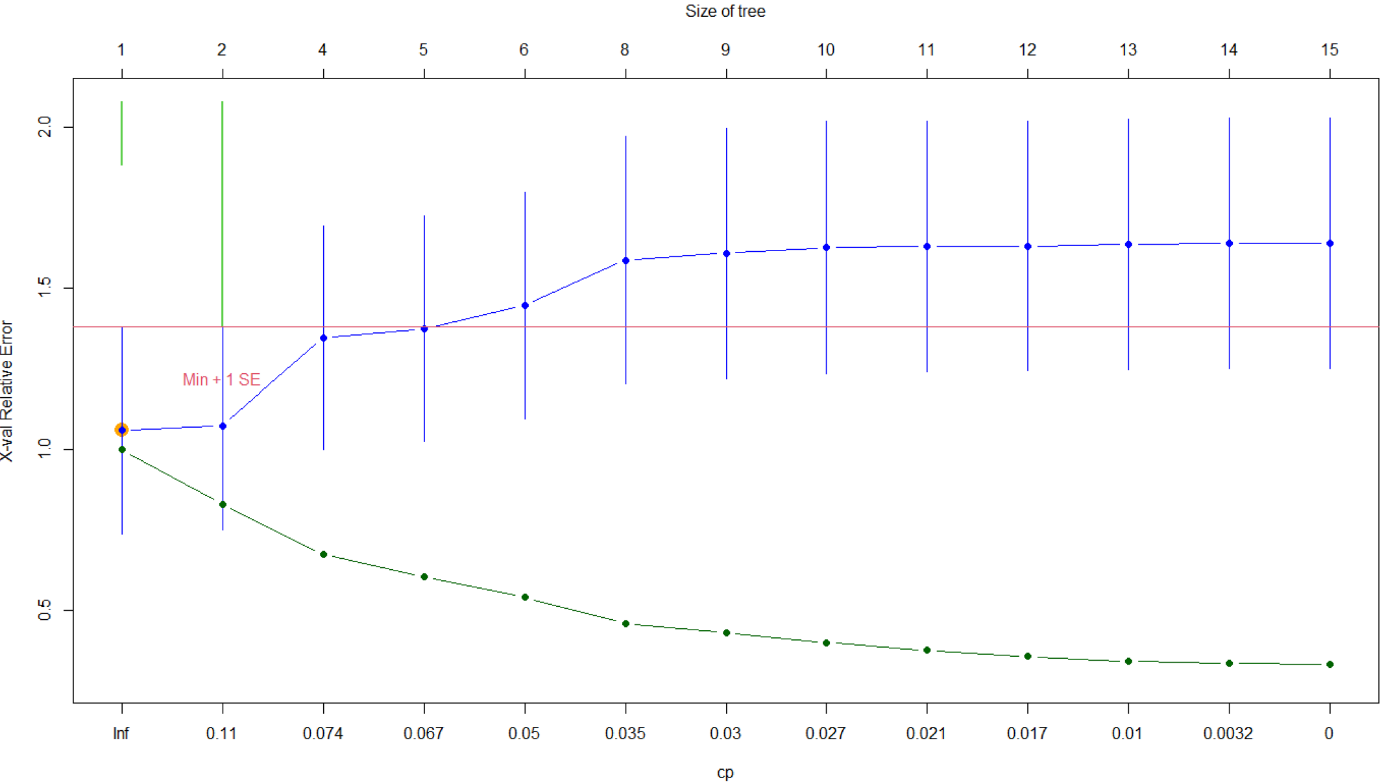

**S3 Fig. Selection of the MRT for the wild hops genetic and metabolic data.** The resulting figure show the relative error (RE) in green, and the cross-validated relative error (CVRE) in blue of trees of increasing size. A CVRE above one indicates that the MRT fails to correctly predict unseen data. The vertical bars indicate one standard error for the CVRE, and the red line indicates one standard error above the minimum CVRE. Orange dot show the smallest tree within one standard error of the CVRE. Lime green bars indicate the number of times each tree size was chosen during the cross-validation process.
