## Supplemental figure 1 for "Fingerprinting and chemotyping approaches reveal a wide genetic and metabolic diversity among wild hops (*Humulus lupulus* L.)"

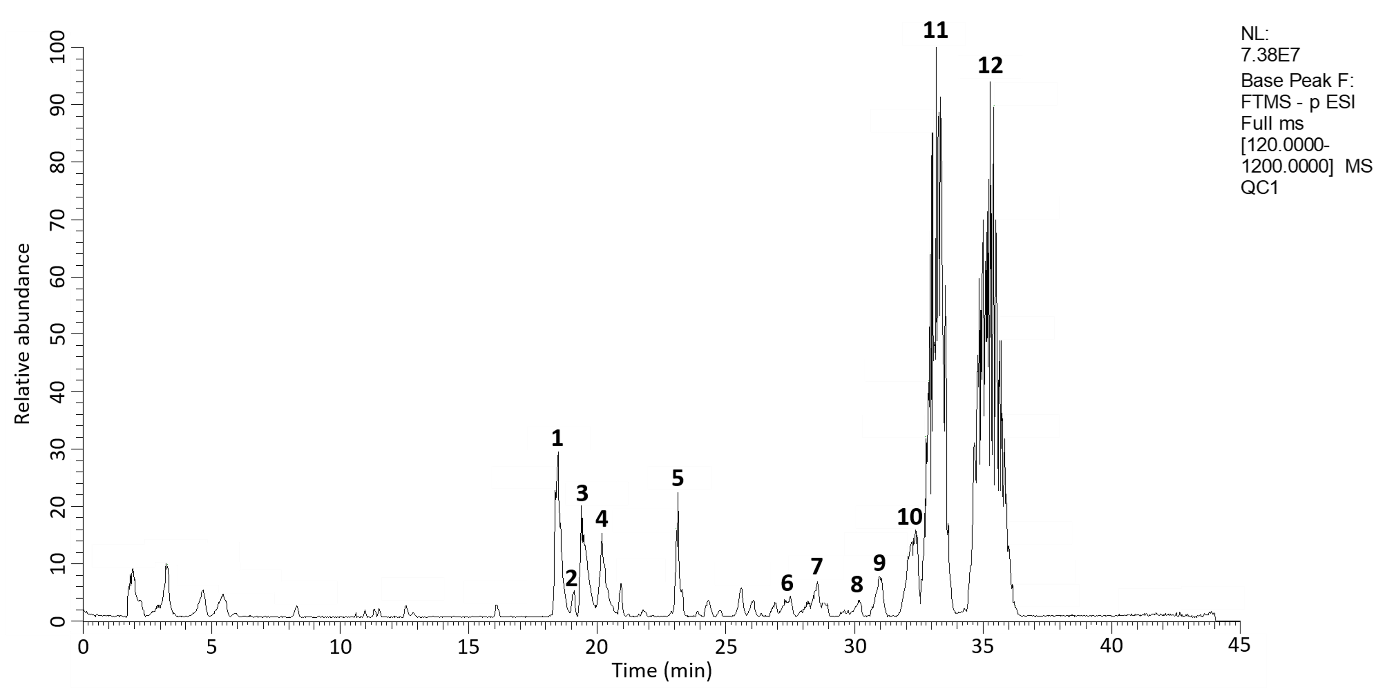

**S1 Fig. UHPLC-MS chromatogram ([M-H]^-^) from Quality Control (QC) of hop leaf extract.** Identified peaks are annotated from 1 to 12 according to S1 Table. NL: Normalization Level, FTMS: Fourier Transform Mass Spectrometry, ESI: Electrospray Ionisation. (XCalibur software - Qual Browser application, Thermo Fisher Scientific).
