## Supplemental figure 2 for "Fingerprinting and chemotyping approaches reveal a wide genetic and metabolic diversity among wild hops (*Humulus lupulus* L.)"

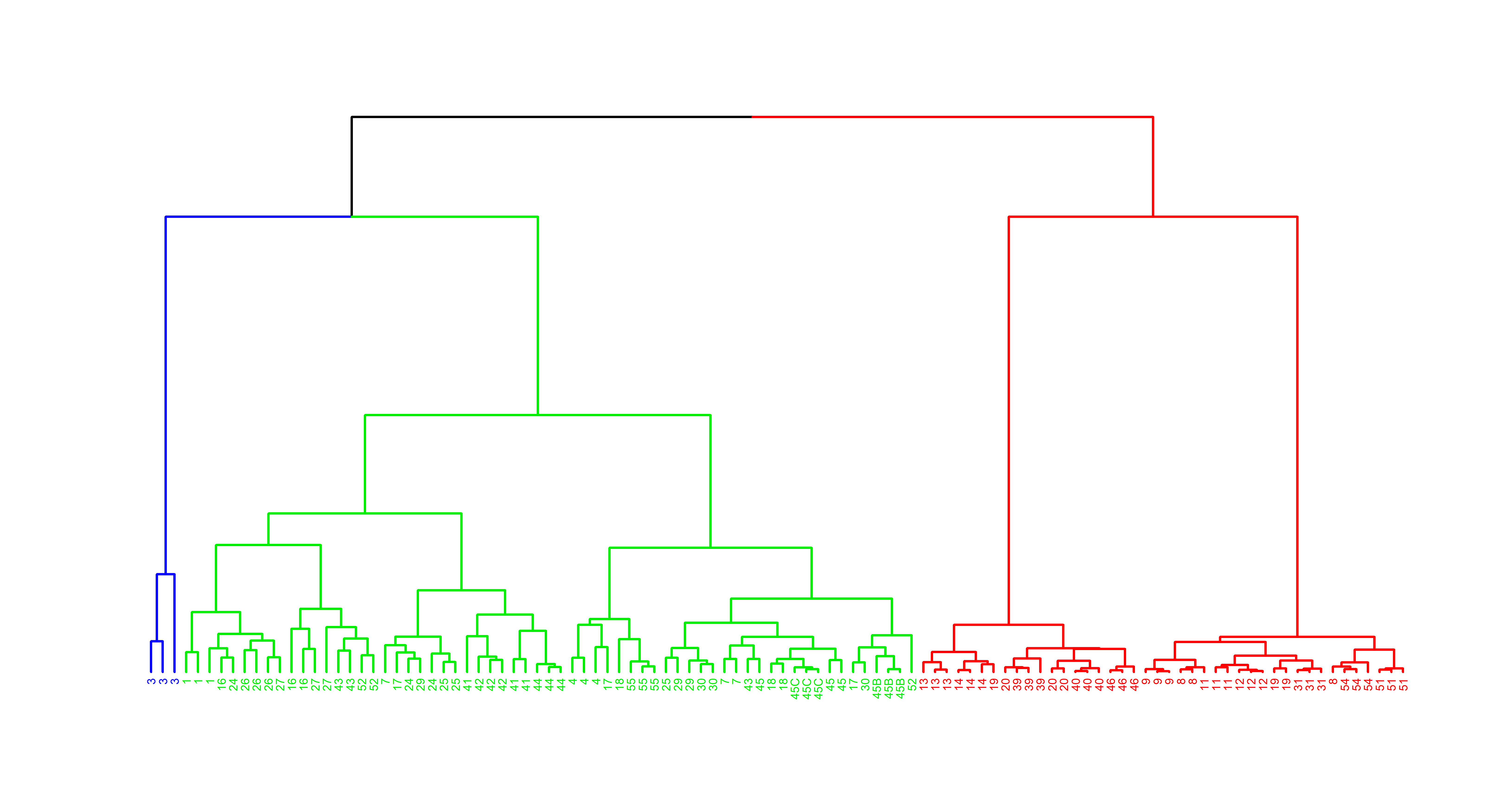

**S2 Fig. Hierarchical Ascending Classification (HAC) analysis based on hops leaf metabolic content.** The resulting dendrogram was made using the “Manhattan” distance, and “ward.D2” clustering method.
